# Novel Immunoglobulin Domain Proteins Provide Insights into Evolution and Pathogenesis Mechanisms of SARS-Related Coronaviruses

**DOI:** 10.1101/2020.03.04.977736

**Authors:** Yongjun Tan, Theresa Schneider, Matthew Leong, L Aravind, Dapeng Zhang

**Affiliations:** Department of Biology, College of Arts and Sciences, Saint Louis University, MO 63110; School of Medicine, Saint Louis University, MO 63110; National Center for Biotechnology Information, National Library of Medicine, National Institutes of Health, Bethesda, MD 20894; Program of Bioinformatics and Computational Biology, College of Arts and Sciences, Saint Louis University, MO 63110

**Keywords:** Coronavirus, COVID-19, SARS, ORF8, Immunoglobin, Evolution, Pathogenicity, Immune evasion

## Abstract

A novel coronavirus (SARS-CoV-2) is the causative agent of an emergent severe respiratory disease (COVID-19) in humans that is threatening to result in a global health crisis. By using genomic, sequence, structural and evolutionary analysis, we show that Alpha- and Beta-CoVs possess several novel families of immunoglobulin (Ig) domain proteins, including ORF8 and ORF7a from SARS-related coronaviruses and two protein groups from certain Alpha-CoVs. Among them, ORF8 is distinguished in being rapidly evolving, possessing a unique insert and a hypervariable position among SARS-CoV-2 genomes in its predicted ligand-binding groove. We also uncover many Ig proteins from several metazoan viruses which are distinct in sequence and structure but share an architecture comparable to that of CoV Ig domain proteins. Hence, we propose that deployment of Ig domain proteins is a widely-used strategy by viruses, and SARS-CoV-2 ORF8 is a potential pathogenicity factor which evolves rapidly to counter the immune response and facilitate the transmission between hosts.

## Introduction

Nidoviruses are an ancient group of lipid-enveloped viruses with non-segmented RNA genomes, which are known to infect oomycetes and animals, including molluscs, arthropods and vertebrates (*1*). Among them are the coronaviruses (CoVs) which possess the largest known monopartite RNA genomes and are classified into four genera—Alphacoronavirus, Betacoronavirus, Gammacoronavirus, and Deltacoronavirus (*2*). Over the past two decades, Beta-CoVs, including the viruses responsible for Severe Acute Respiratory Syndrome (SARS) in 2003 and Middle Eastern Respiratory Syndrome (MERS) in 2012, have emerged as significant local land global health concerns with economic consequences (*3, 4*). Recently, a novel severe respiratory disease has emerged in humans (abbreviated COVID-19) (*5, 6*). COVID-19 presents with a relatively long incubation period of 1-2 weeks followed by development of fever, dry cough, dyspnea and bilateral ground-glass opacities in the lungs (*7*). In some patients, this can proceed to fatal respiratory failure, characterized by acute lung injury (*8*) and acute respiratory distress syndrome (*9*). Within several months of the first outbreak, there have been over 75,000 confirmed cases with over 2,000 deaths of COVID-19 globally (https://www.who.int/docs/default-source/coronaviruse/situation-reports/20200220-sitrep-31-covid-19.pdf). Due to the rapid spread and potential severity of the disease, COVID-19 poses as a potential major threat to human health. A novel coronavirus (SARS-CoV-2) was identified as the causative agent of COVID-19, and phylogenomic analysis has shown that it belongs to the same larger clade of Beta-CoVs as the SARS-CoV with a likely origin in bats (*5, 10, 11*). Despite intense scrutiny, multiple proteins encoded by the SARS-related CoV (including SARS-CoV-2) genome remain enigmatic. Here, we present a computational and evolutionary analysis to show that one such mysterious protein, ORF8, and several others from Alpha- and Beta-coronavirus, comprise novel families of immunoglobulin domain proteins, which might function as potential modulators of host immunity to delay or attenuate the immune response against the viruses.

## Materials and methods

### Genome comparison analysis

We retrieved the SARS-related CoV genomes by searching against the non-redundant (nr) nucleotide database of the National Center for Biotechnology Information (NCBI) with the SARS-CoV-2 genome sequence (NC_045512.2) as a query (*12*). The program CD-HIT was used for similarity-based clustering (*13*). A multiple sequence alignment of whole virus genomes was performed by KALIGN (*14*). Based on the MSA, a similarity plot was constructed by a custom Python script, which calculated the identity between each subject sequence and the SARS-CoV-2 genome sequence based on a custom sliding window size and step size. Open reading frames of virus genomes used in this study were extracted from an NCBI Genbank file.

### Protein sequence analysis

To collect protein homologs, iterative sequence profile searches were conducted by the programs PSI-BLAST (Position-Specific Iterated BLAST)(*12*) and JACKHMMER (*15*), which searched against the non-redundant (nr) protein database of NCBI with a cut-off e-value of 0.005 serving as the significance threshold. Similarity-based clustering was conducted by BLASTCLUST, a BLAST score-based single-linkage clustering method (ftp://ftp.ncbi.nih.gov/blast/documents/blastclust.html). Multiple sequence alignments were built by the KALIGN (*14*), MUSCLE(*16*) and PROMALS3D(*17*) programs, followed by careful manual adjustments based on the profile–profile alignment, the secondary structure information and the structural alignment. Profile-profile comparison was conducted using the HHpred program (*18*). The consensus of the alignment was calculated using a custom Perl script. The alignments were colored using an in-house alignment visualization program written in perl and further modified using adobe illustrator. Signal peptides were predicted by the SignalP-5.0 Server (*19*). The transmembrane regions were predicted by the TMHMM Server v. 2.0 (*20*).

### Identification of distinct viral Ig domain proteins

By using the protein remote relationship detection methods, we generated a collection of distinct Ig domains from the Pfam database (*21*) and also from our local domain database. Then, we utilized the hmmscan program of the HMMER package (*22*) and RPS-BLAST (*12, 23*) to retrieve the homologs from viral genomes.

### Molecular Phylogenetic analysis

The evolutionary history was inferred by using the Maximum Likelihood method based on the JTT w/freq. model (*24*). The tree with the highest log likelihood is shown. Support values out of 100 bootstraps are shown next to the branches (*25*). Initial tree(s) for the heuristic search were obtained automatically by applying Neighbor-Join and BioNJ algorithms to a matrix of pairwise distances estimated using a JTT model, and then selecting the topology with the superior log likelihood value. A discrete Gamma distribution was used to model evolutionary rate differences among sites (4 categories). The rate variation model allowed for some sites to be evolutionarily invariable. The tree is drawn to scale, with branch lengths measured in the number of substitutions per site. The tree diagram was generated using MEGA Tree Explorer (*26*)

### Entropy analysis

Position-wise Shannon entropy (H) for a given multiple sequence alignment was calculated using the equation:

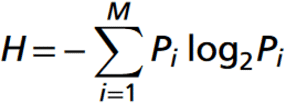

*P* is the fraction of residues of amino acid type *i*, and *M* is the number of amino acid types. The Shannon entropy for the *i*th position in the alignment ranges from 0 (only one residue at that position) to 4.32 (all 20 residues equally represented at that position). Analysis of the entropy values which were thus derived was performed using the R language.

### Protein Structure Prediction and Analysis

The secondary structural prediction was conducted using the Jnet (Joint Network) program (*27*). Jnet is a neural network-based predictor which trains neural networks from three different types of profiles: profile PSSM, profile HMM, and residue frequency profile. It generates a consensus secondary structure with an average accuracy of 72% or greater.

The Modeller9v1 program (Sali and Blundell, 1993) was utilized for homology modeling of structure of SARS-CoV-2 ORF8 using the SARS-CoV ORF7 (1xak_A) (*28*) as a template. Since in these low sequence-identity cases, sequence alignment is the most important factor affecting the quality of the model (Cozzetto and Tramontano, 2005), alignments used in this study have been carefully built and cross-validated based on the information from HHpred and edited manually using the secondary structure information. We generated five models and selected the one that had better model accuracy p-value and global model quality score as assessed by ModFOLD6 online server (*29*). Structural analysis and comparison were conducted using the molecular visualization program PyMOL (*30*). The structural similarity search was performed using the DALI server (*31*).

## Results and discussion

### Comparative genomics unveils fast-evolving regions of SARS-related CoV genomes

The host-pathogen arms-race has selected for a disparate complement of viral genes involved in pathogenesis. These genes often rapidly diversify through recombination and mutations to keep up with the evolution of host resistance. To identify proteins with potential pathogenic roles in COVID-19, we conducted a comparative genomic analysis of the coronaviruses. Similarity plots show that the bat CoV RaTG13 is the closest relative of SARS-CoV-2 with no evidence for recombination between them (Figure 1 and Figure S1). SARS-CoV-2 also shows good similarity to two other bat viruses, CoVZXC21 and CoVZXC45, first in the 5’ half of ORF1 and again after nucleotide number 20,000 of the genome. However, the remaining part of ORF1 of SARS-CoV-2 and RatG13 show no specific relationship to these viruses. This suggests a recombination event between the common ancestor of SARS-CoV-2 and RaTG13, and probably another member of the SARS-related clade close to CoVZXC21 and CoVZXC45. In addition to this major recombination event, we identified several smaller regions which might have undergone recombinational diversification during the emergence of the SARS-CoV-2 genome (Figure S1). Notably, many of them are clustered in three regions displaying extensive diversification, corresponding to the N-terminal region of the ORF1a polyprotein, the Spike protein, and the uncharacterized protein encoded by ORF8 (Figure 1 and Figure S1).

**Figure 1.**
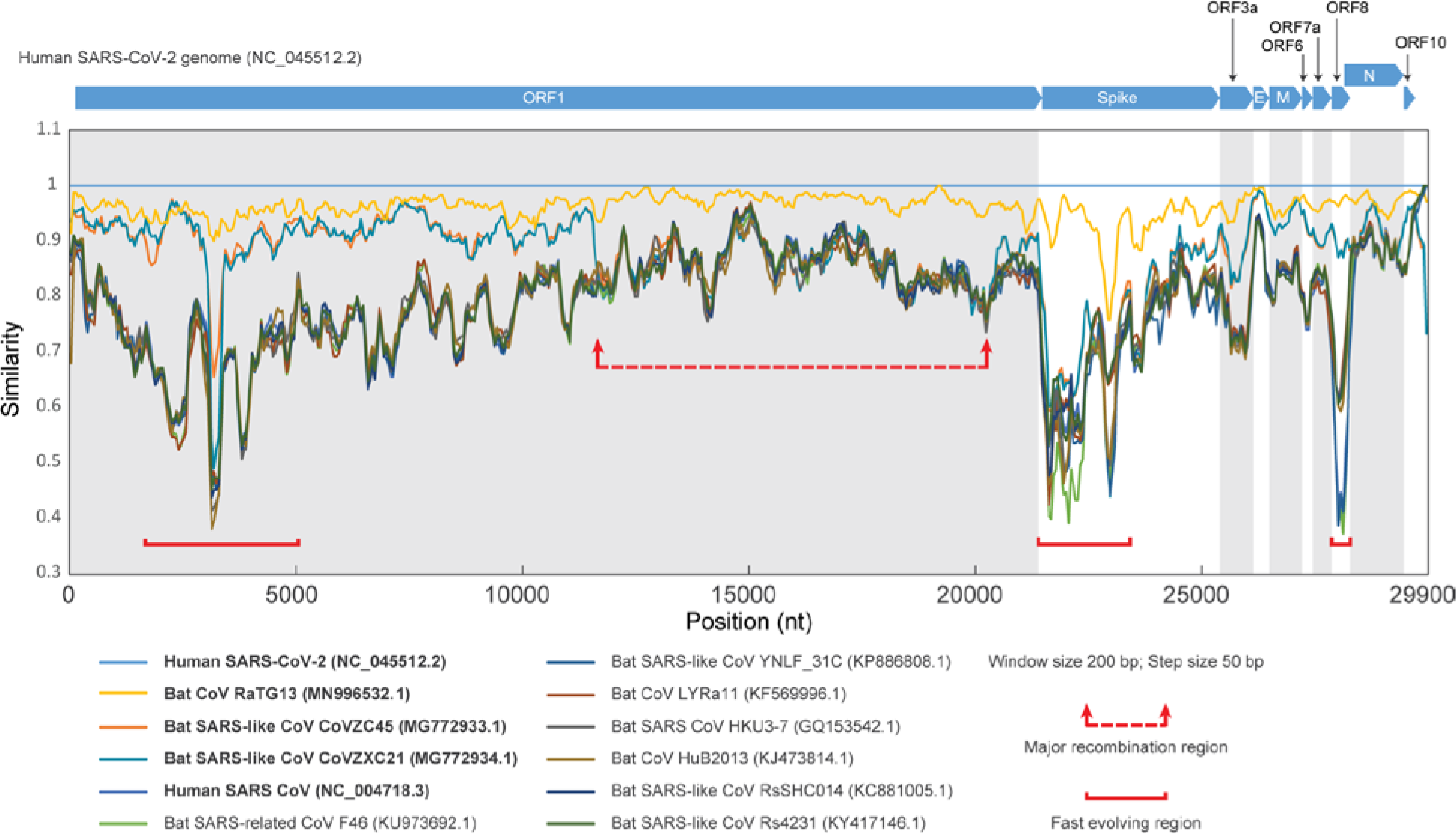
Genome comparison analysis of SARS-related CoVs. Similarity plot of SARS-related CoVs against human SARS-CoV-2 Wuhan-Hu-1 genome (NC_045512.2), based on a multiple sequence alignment of the whole genomes. Each point represents the percent identity of a 200 bp window of the alignment with a 50 bp step size between each point. The open reading frames of the SARS-CoV-2 genome (NC_045512.2) are shown above the plot. Each colored line corresponds to the nucleotide similarity between the human SARS-CoV-2 genome (NC_045512.2) and the respective SARS-related CoV genome. The red arrows and dashed line surround a region displaying major divergence due to possible recombination within SARS-related CoV genomes. The regions marked by a solid red line highlight fast-evolving regions among the SARS-related CoV genomes. For detailed information about the genomes that were used in this study, refer to Table S1.

### Identification of novel immunoglobin protein families in CoVs

Among these three fast-evolving proteins, we focused on the ORF8 protein as it is one of the so-called accessory proteins, which does not participate in viral replication (*32, 33*), raising the possibility that it might have a role in viral pathogenesis. It is a predicted secreted protein present only in some Beta-CoVs, including SARS-CoV-2 but not the MERS-like clade. Profile-profile comparisons using a sequence-profile built from the multiple sequence alignment of all available ORF8 proteins showed it to be unexpectedly homologous to the membrane-anchored ORF7a protein from the same subset of Beta-CoVs, and several proteins (variously annotated as ORF9 or ORF10) from a subset of bat Alpha-CoVs (Figure 2A) (probability=94% of profile-profile match) (*34*). ORF7a is a known member of the immunoglobulin (Ig) domain superfamily and is specifically related to extracellular metazoan Ig domains that are involved in adhesion, such as ICAM (*35, 36*). The Beta-CoV ORF8, ORF7a and the Alpha-CoV Ig domains display a classic β-sandwich fold with seven β-stands and share the characteristic pattern of two cysteines which form stabilizing disulfide bonds with metazoan Ig domains (Figure 2A and Figure S2) (*37*). However, they are unified as a clade by the presence of an additional pair of conserved disulfide-bonding cysteines (Figure 2A). Nonetheless, there are notable structural differences between the three groups of proteins. ORF8 is distinguished from ORF7a and Alpha-CoV Ig proteins by the loss of the C-terminal transmembrane (TM) helix and the acquisition of a long insert between stands 3 and 4 with a conserved cysteine which might facilitate dimerization through disulfide-bond formation (Figure 2A). The homology model of ORF8, based on the structure of SARS-CoV ORF7a (pdb id:1XAK), suggests that this insert augments a potential peptide-ligand binding groove that has been proposed for ORF7a (Figure 2B and Figure S3). Hence, the emergence of the insert has gone hand-in-hand with the acquisition of a modified interaction interface.

**Figure 2.**
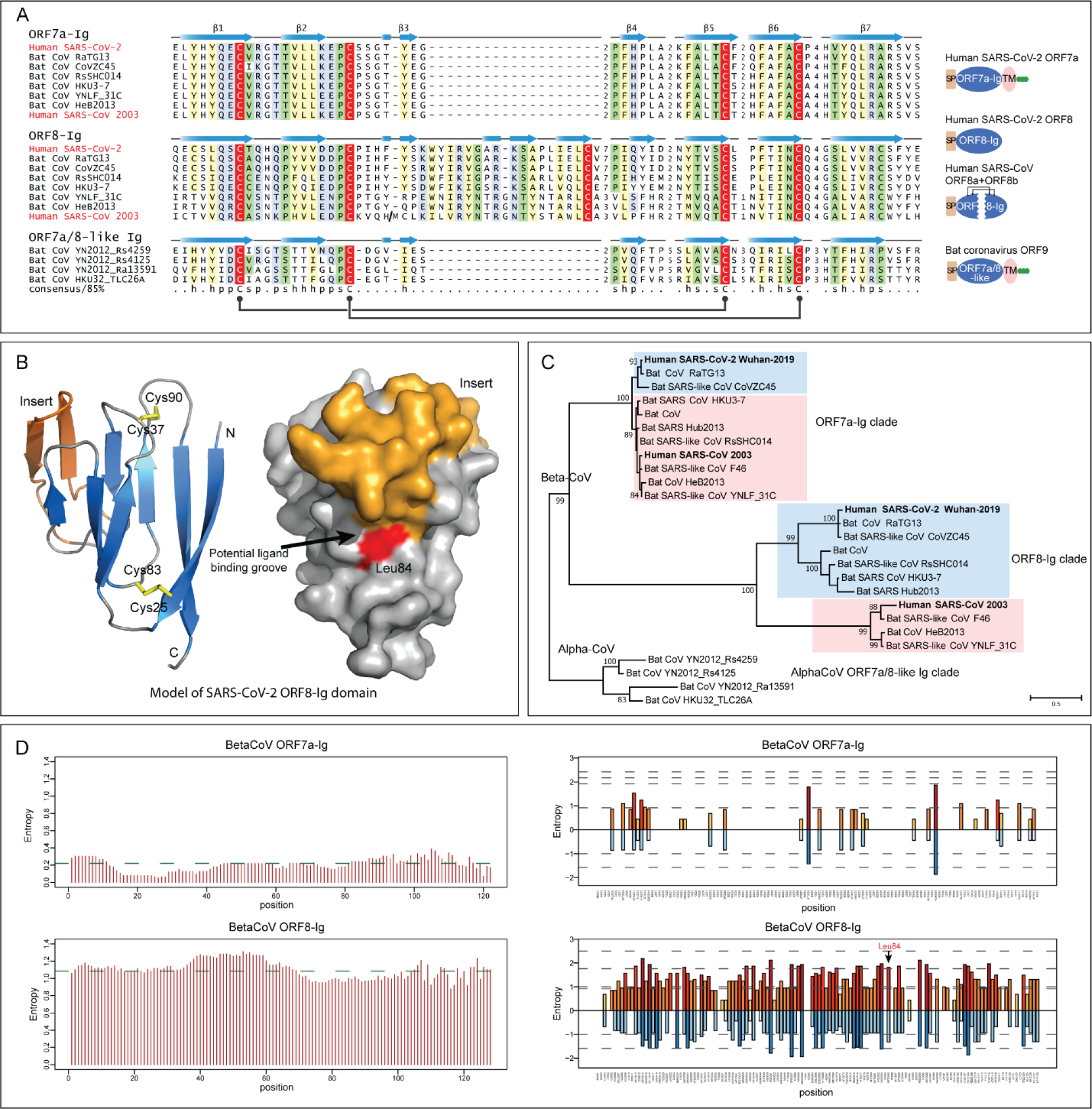
Sequence, structure and evolutionary analysis of novel Ig domain proteins in SARS-related CoVs. (A) Multiple sequence alignment (MSA) and representative domain architectures of ORF7a-Ig, ORF8-Ig, and ORF7a/8-like Ig domain families. Each sequence in the MSA was labelled by its species abbreviation followed by its source. The predicted secondary structure is shown above each alignment and the consensus is shown below the super-alignment, where h stands for hydrophobic residues, s for small residues, and p for polar residues. Two pairs of conserved cysteines that form disulfide bonds are highlighted in red. (B) Homology model of SARS-CoV-2 ORF8-Ig domain (YP_009724396.1) and the location of the hypervariable position corresponding to Leu84 in the predicted ligand-binding groove. The β-sheets of the common core of the Ig fold are colored in blue, the insert in ORF8-Ig in orange and the loops in grey. The characteristic disulfide bonds are highlighted in yellow. (C) Maximum likelihood phylogenetic analysis of CoV Ig domain families. Support values out of 100 bootstraps are shown for the major branches only. (D) Entropy plot for the ORF7a and ORF8 proteins in betacoronavirus. Left: Shannon entropy computed for each column for a character space of 20 amino acids and presented as mean entropy in a sliding window of 30 residues. The mean entropy across the entire length of the protein is indicated as a green horizontal line. Right: Shannon entropy in regular amino acid alphabet (20 amino acids) are shown above the zero line in shades of orange. Shannon entropy in a reduced alphabet of 8 residues are shown below the zero line in shades of blue. If a position shows high entropy in both alphabets it is a sign of potential positive selection at those positions for amino acids of different chemical character.

Besides these families, we identified a fourth family of Ig domains from the same Alpha-CoVs which contain the above-discussed ORF7a/8-like Ig family (Figure 3A). These Alpha-CoVs typically possess one or two paralogous copies annotated as either ORF4a/b or NS5a/b. From their sequences, these Ig domains are not closely related to the ORF7a and ORF8 Ig domains (Figure S4). However, profile-profile searches have shown that they are related to Ig domains found in the adenoviral E3-CR1 proteins (probability: 90% of matching the Pfam CR1 Ig domain profile) (Figure S4). In these searches, they also yield weaker hits to two other Ig domains, namely the poxviral decoy interferon receptors and human T-cell surface CD3 zeta (Figure S4).

**Figure 3.**
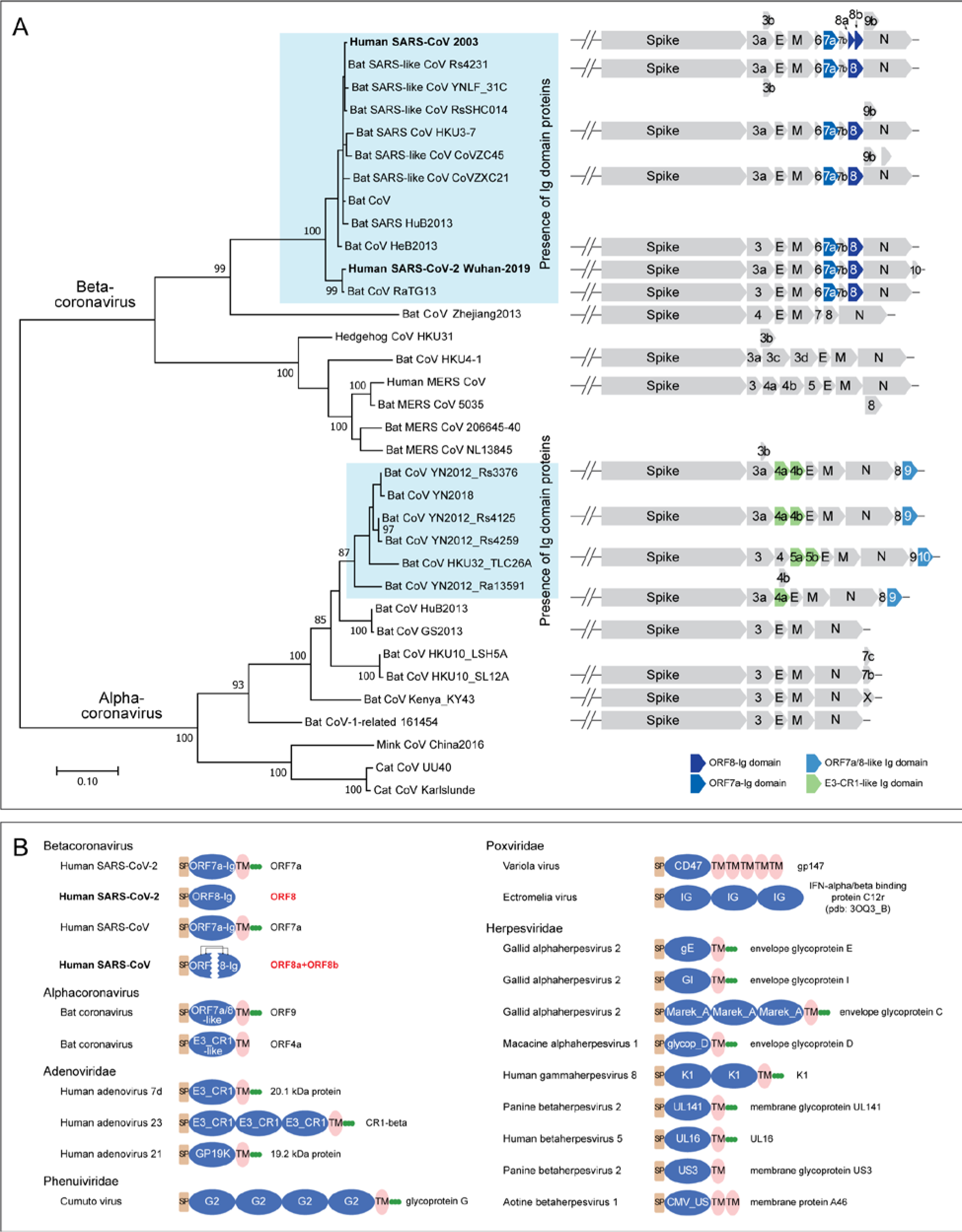
(A) Genomic structure analysis of SARS-related CoVs. The tree of coronavirus was built based on an MSA of a coronavirus RNA-directed, RNA polymerase domain using a maximum likelihood model. Supporting values from 100 bootstraps are shown for the major branches only. The genome structure of major representative CoVs are shown right to the terminal clade of the phylogenetic tree. (B) Representative domain architectures of the Ig components in different animal viruses. Proteins were grouped based on their families, except for proteins of coronavirus, which were grouped based on their genus. For the information of the NCBI accession numbers, refer to the supplementary data.

### ORF8 is a fast-evolving protein in SARS-related CoVs

Phylogenetic analysis of ORF7a, ORF8 and Alpha-CoV Ig domains shows that each group represents a distinct clade (Figure 2C). The tree topology of ORF7a mirrors that of the polymerase tree (Figure 3A); however, the topology of the ORF8-Ig clade is not consistent with it. This might be due to a recombination event between the SARS-related CoVs (as suggested by the similarity plot analysis) and/or unusual divergence under selection. To better understand the functional difference between ORF8 and ORF7a, we examined the column-wise Shannon entropy in the 20 amino acid alphabet and found that the ORF8 has significantly higher mean entropy than ORF7a (ORF8: 1.09 vs ORF7a: 0.22, p<10^−16^ for the H_0_ of congruent means by t-test) (Figure 2D). By comparing column-wise entropies in both the 20 amino acid and a reduced 8-letter alphabet (where amino acids are grouped based on similar side-chain chemistry), we found at least 14 positions in ORF8 which show high entropy in both alphabets as compared to a single position in ORF7a (Figure 2D). This indicates that ORF8 is a fast-evolving protein under selection for diversity as contrast to ORF7a. Strikingly, one of these highly variable positions, which features residues with very different side characters (hydrophobic, acidic, alcoholic and proline), corresponding to Leu84 was also identified as the most variable position across 54 closely related human SARS-CoV-2 genome sequences (*38*). In our structural model, this residue is positioned at the predicted peptide-ligand binding groove of the ORF8-Ig domain (Figure 2B). Therefore, our entropy and structural analysis of the ORF8-Ig domain, in conjunction with its hypervariable position found in human SARS-CoV-2 genomes, points to a role for ORF8 at the interface of the host-virus interaction possibly in a pathogenic context.

### Ig domain proteins are newly acquired in subsets of Alpha- and Beta-CoVs

We examined the distribution of CoV Ig proteins in the context of a phylogenetic tree of both Beta-and Alpha-CoVs based on their polymerase proteins (Figure 3A). Other than the two subsets of Beta-CoVs and Alpha-CoVs that contain the above-described Ig domain proteins, no other CoVs contain any Ig domain proteins (Figure 3A). The immediate sister-groups of the Ig-containing CoVs typically have Spike, E, M and N, and one or two other uncharacterized accessory proteins which are not Ig domains to our best knowledge. The Alpha-CoV ORF9/10 share a C-terminal TM helix and, along with ORF7a of the Beta-CoVs, lack the insert in the Ig domain (Figure 2A). Hence, it is possible that this architecture represents the ancestral state which was present in the common ancestor of both Alpha-CoVs and Beta-CoVs. Under this scenario, the protein was displaced/lost both in certain Alpha-CoVs and Beta-CoVs. Alternatively, ORF7a could have been exchanged between Alpha- and Beta-CoVs by a recombination event. In both scenarios, ORF8 arose likely via a duplication of ORF7a in specifically the Beta-CoVs. Although we couldn’t identify the ultimate precursors of the CoV Ig domains, they are likely to have been acquired on at least two independent occasions from different sources. The CoV ORF7a-ORF8 families might have derived from the metazoan adhesion Ig families, and the ORF4a/b-like Ig domains of Alpha-CoVs were likely acquired from adenoviral CR1 Ig domains with which they share some specific sequence features.

### Divergent Ig proteins with a comparable architecture are deployed by distinct viruses

The presence of multiple Ig domains with different affinities in CoVs prompted us to more generally survey animal viruses for Ig domains. By using the Pfam models (*39*) and our own PSSMs created from PSI-BLAST runs with Ig domains (*12*), we were able to identify about 17 distinct viral Ig domain families in a wider diversity of animal viruses (Figure 3B and Table S2). In addition to CoVs, such Ig domain proteins can be found in adenoviruses, NCLDVs, Herpesviruses and Phenuiviruses. These viral Ig domains are highly divergent; many of them are only found in certain viral groups. However, the majority have an architecture comparable to the CoV-Ig domains, with an N-terminal signal peptide, one or multiple Ig domains and a C-terminal TM region often followed by a stretch of basic residues. Thus, although the Ig domains are not the universally preserved component of viruses, they have been acquired and selected independently by a wide range of viruses. The presence of a proofreading 3’-5’ exonuclease has been proposed to favor the emergence of larger RNA genomes in CoVs (*40*). Indeed, this might have also contributed to the acquisition of potential pathogenesis factors such as the Ig domains described herein which are comparable to those seen in DNA viruses.

### Novel CoV Ig domain proteins are potential immune modulators

Why have diverse viruses independently acquired the Ig domain during their evolution? First, the Ig domains are major mediators of adhesive interactions in both eukaryotes and prokaryotes (*37, 41, 42*). Thus, this domain can be used for adherence for cell to cell spread (e.g. herpesviral Ig domain proteins) (*43*). Further, Ig domains are major building blocks of metazoan immune systems. Thus, viruses often utlize this domain to disrupt immune signaling of the host. For example, in adenoviruses, the CR1 Ig domain proteins have been shown to block the surface expression on infected cells of class I major histocompatibility complex molecules by blocking their trafficking from the endoplasmic reticulum (ER) to Golgi (*44*). This has been shown to affect the host inflammatory response and modulates the presentation of viral antigens to T-cells (*45*). In poxvirus, the secreted Ig domain proteins function as interferon receptors or decoys that bind the interferon-α/β and disrupt signaling via the endogenous host receptors (*46*). Further, SARS-ORF7a has been implicated in the interaction with bone marrow stromal antigen 2 (BST-2), which tethers budding virions to the host cell in a broad-spectrum antiviral response, to prevent the N-linked glycosylation of BST-2 thereby crippling the host response against the virus (*47*). Given their shared evolutionary history and similar sequence and structural features, we predict that the newly identified CoV Ig domain proteins, such as ORF8 of SARS-CoV-2, might similarly function as immune modulators.

While ORF8 is a paralog of ORF7a, its lack of the TM segment, unique insert and significantly more rapid evolution suggest that it has acquired a distinct function and has been under strong positive selection One possible mechanism is that, like the adenoviral CR1 proteins, it interferes with MHC molecules to attenuate antigen presentation, resulting in ineffective detection of the virus by the host immune system. Consistent with this, the SARS-CoV ortholog translocates to the ER (*48*) and its higher variability indicates probable selection due to its interaction with a rapidly evolving host molecule. Notably, while the SARS-CoV ORF8 isolates from civets and early stages of the human epidemic are intact, it split up into two ORFs (ORF8a and ORF8b) during the subsequent human epidemic (*49*). ORF8a and ORF8b retain the conserved Cys residues of the Ig domain and have been observed to form a complex in a yeast-two hybrid interaction study (*50*). This suggests it might still fold into an intact structure held by the two disulfide bonds formed by four conserved Cys residues.

In conclusion, the presence of fast-evolving ORF8 Ig domain proteins in the SARS-related viruses, including the emergent 2019 SARS-CoV-2, suggests that they might be potential pathogenicity factors which counter or attenuate the host immune response and might have facilitated the transmission between hosts. We hope that the discovery and analyses of the novel Ig domain proteins reported here will help the community better understand the evolution and pathogenesis mechanisms of these coronaviruses.

## Acknowledgments

Y.T., T. S., M.L., and D. Z. were supported by Saint Louis University start-up fund. L.A. was supported by the Intramural Research Program of the NIH, National Library of Medicine.

## SUPPLEMENTAL INFORMATION

**Supplementary Figure S1.**
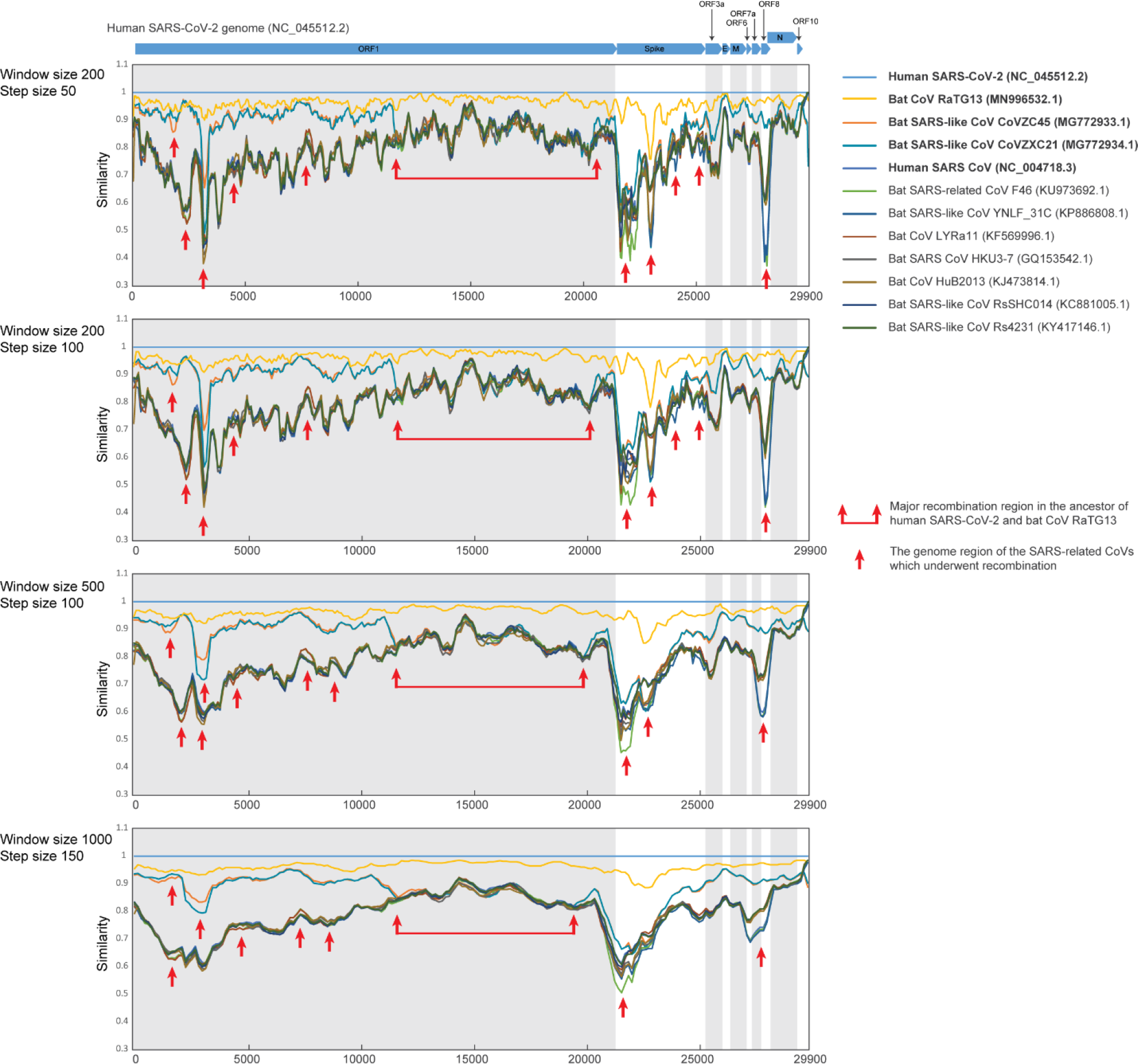
Genome comparison analysis of SARS-related CoVs. Similarity Plot of SARS-related CoVs against human SARS-CoV-2 Wuhan-Hu-1 genome (NC_045512.2) based on a multiple sequence alignment of the whole genomes. Each point represents a different slicing window size from the alignment with a different step size between each point. For each plot, the window size and step size are shown in the top left. Horizontal bars above the top plot correspond to the different open reading frames of the SARS-CoV-2 genome (NC_045512.2). Each different colored line corresponds to the nucleotide similarity between the human SARS-CoV-2 genome (NC_045512.2) and the respective SARS-related CoV genome. The red arrows and solid lines surround regions which display recombination within the SARS-related CoV genomes. The single red arrows point to specific regions of recombination.

**Supplementary Figure S2.**
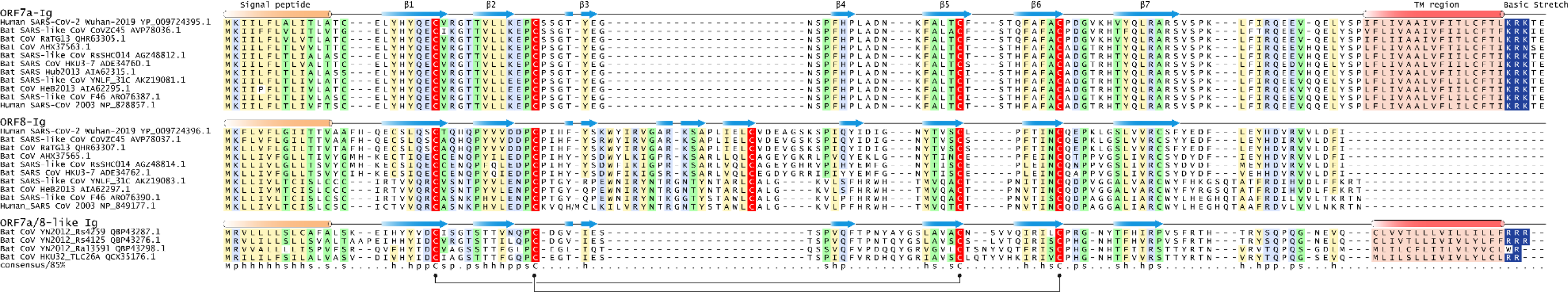
Full length multiple sequence alignment of ORF7a, ORF8-Ig and ORF7a/8-like proteins. Each sequence in the MSA was labelled by its species abbreviation followed by its isolation and NCBI accession number. The predicted secondary structure is shown above the alignment and the consensus is shown below the alignment, where h stands for hydrophobic residues, s for small residues, and p for polar residues. The characteristic signal peptide, TM region and a stretch of basic residues are also labeled.

**Supplementary Figure S3.**
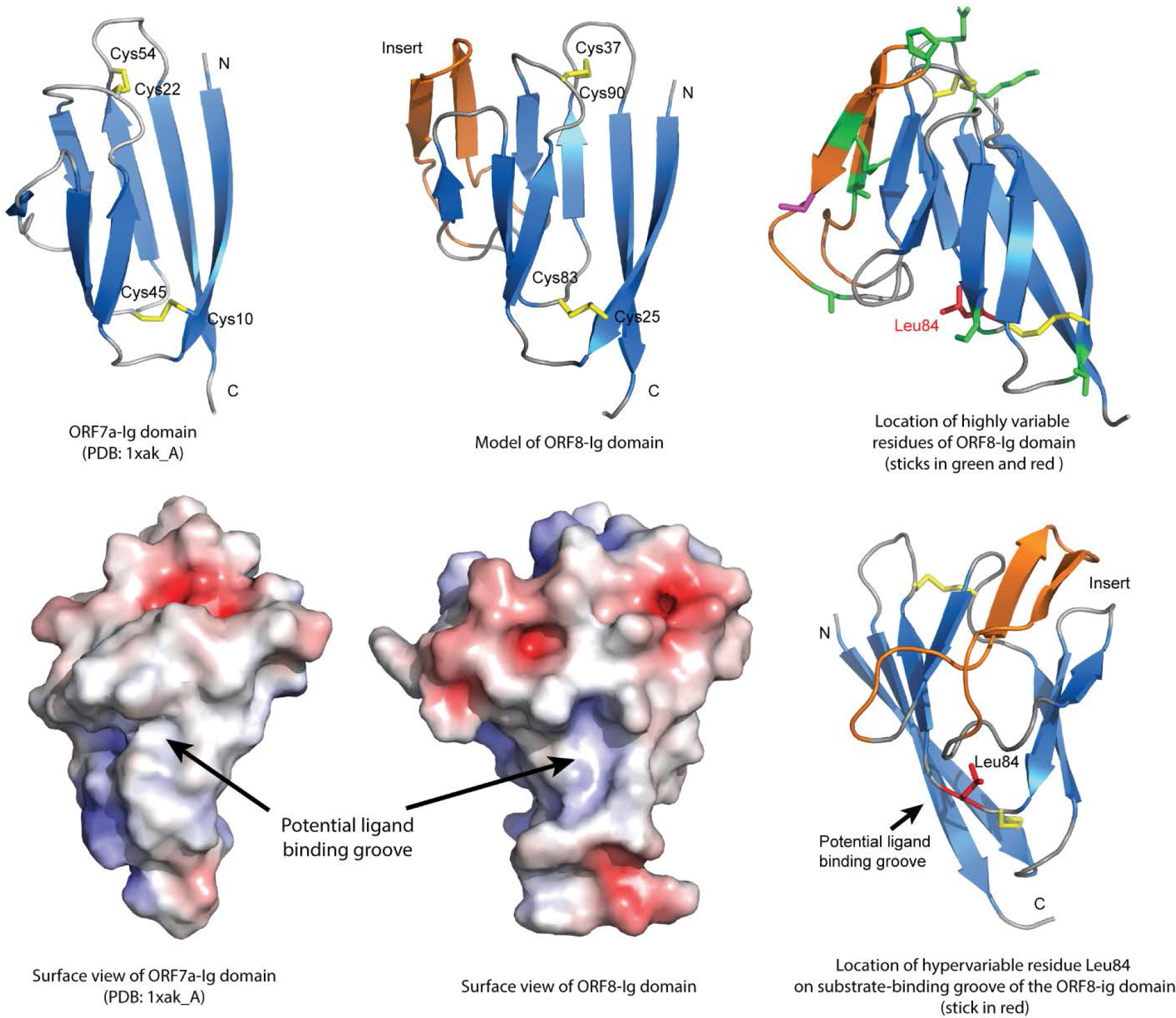
Structural analysis of CoV Ig domains.

**Supplementary Figure S4.**
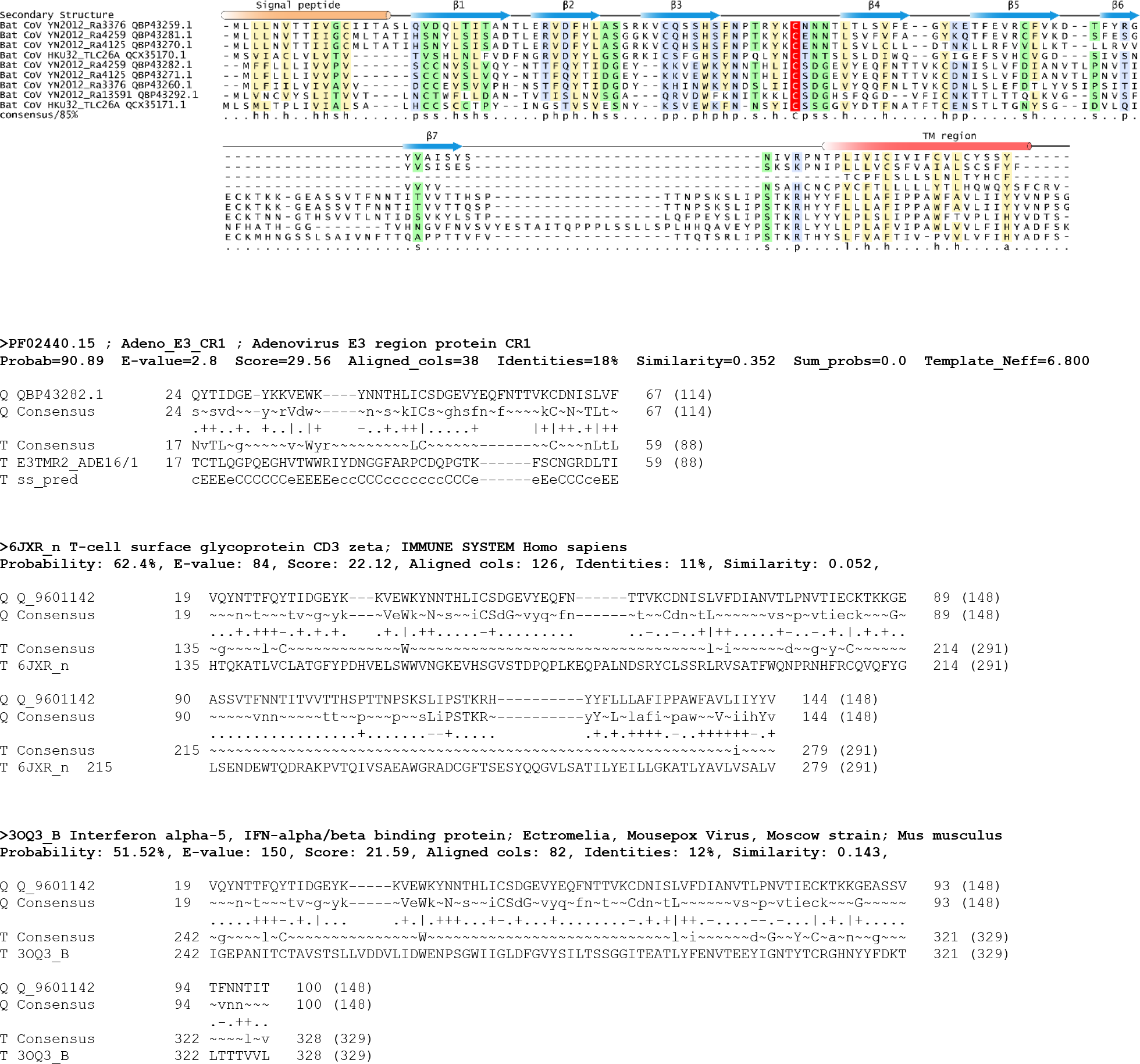
Multiple sequence alignment of alphacoronavirus E3-CR1-like Ig domain proteins and their related Ig domains identified by profile-profile searches.

**Supplementary Table S1.** Detailed information of Human SARS-CoV-2 Wuhan-Hu-1 genome and other SARS-related genomes that were used in this study (Figure 1 & Figure S1).

| Organism | Host | NCBI ID | Year | Citation |
| --- | --- | --- | --- | --- |
| Severe acute respiratory syndrome coronavirus 2 (SARS-CoV-2) isolate Wuhan-Hu-1 | Human | NC_045512.2 | 2020 | A novel coronavirus associated with a respiratory disease in Wuhan of Hubei province, China (Unpublished) |
| Bat SARS-like coronavirus isolate bat-SL-CoVZC45 | Bat | MG772933.1 | 2018 | (1) |
| Bat SARS-like coronavirus isolate bat-SL-CoVZXC21 | Bat | MG772934.1 | 2018 | (1) |
| Bat coronavirus isolate RaTG13 | Bat | MN996532.1 | 2020 | Not Available |
| Severe acute respiratory syndrome-related coronavirus | Human | NC_004718.3 | 2003 | (2) |
| Severe acute respiratory syndrome-related coronavirus isolate F46 | Bat | KU973692.1 | 2017 | Identification of a new intermediate virus between bat-CoVs and SARS-CoVs from least horseshoe bats in China (Unpublished) |
| Bat SARS-like coronavirus YNLF_31C | Bat | KP886808.1 | 2015 | Not Available |
| Rhinolophus affinis coronavirus isolate LYRa11 | Bat | KF569996.1 | 2014 | (3) |
| Bat SARS coronavirus HKU3-7 | Bat | GQ153542.1 | 2010 | (4) |
| BtRs-BetaCoV/HuB2013 | Bat | KJ473814.1 | 2015 | (5) |
| Bat SARS-like coronavirus RsSHC014 | Bat | KC881005.1 | 2013 | (6) |
| Bat SARS-like coronavirus isolate Rs4231 | Bat | KY417146.1 | 2017 | (7) |

**Supplementary Table S2.**
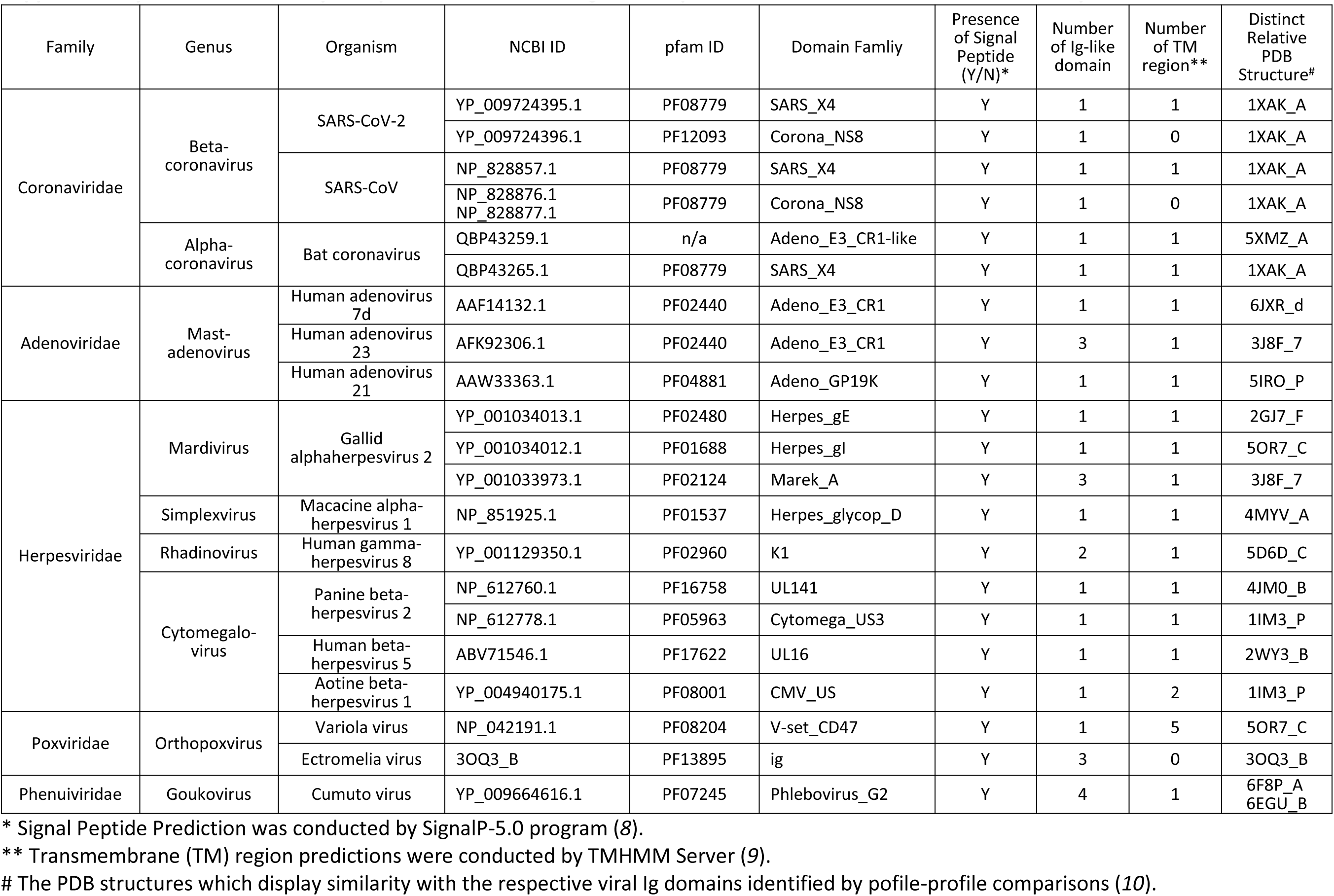
Summary of representatives of viral Ig domain proteins which were identified in this study.

